# A gustatory receptor tuned to the steroid plant hormone brassinolide in *Plutella xylostella* (Lepidoptera: Plutellidae)

**DOI:** 10.1101/2020.07.13.200048

**Authors:** Ke Yang, Xin-Lin Gong, Guo-Cheng Li, Ling-Qiao Huang, Chao Ning, Chen-Zhu Wang

## Abstract

Feeding and oviposition deterrents help phytophagous insects to identify host plants. The taste organs of phytophagous insects contain bitter gustatory receptors (GRs). To explore their function, we focused on PxylGr34, a bitter GR in *Plutella xylostella* (L.). We detected abundant *PxylGr34* transcripts in the larval head and specific expression of *PxylGr34* in the antennae of females. Analyses using the *Xenopus* oocyte expression system and two-electrode voltage-clamp recording showed that PxylGr34 is specifically tuned to the plant hormone brassinolide (BL) and its analog 24-epibrassinolide. Electrophysiological analyses revealed that the medial sensilla styloconica on the maxillary galea of the 4^th^ instar larvae are responsive to BL. Dual-choice bioassays demonstrated that BL inhibits larval feeding and female ovipositing. Knock-down of PxylGr34 by RNAi abolished BL-induced feeding inhibition. These results shed light on gustatory coding mechanisms and deterrence of insect feeding and ovipositing, and may be useful for designing plant hormone-based pest management strategies.

## Introduction

Many phytophagous insects have evolved to select a limited range of host plants. Understanding the ultimate and proximate mechanisms underlying this selection strategy is a crucial issue in the field of insect–plant interactions. Insects’ decisions to feed and oviposit are mainly based on information carried via chemosensory systems (Schoonhoven et al., 2005). Chemosensory perception of feeding/oviposition deterrents by phytophagous insects plays an important role in host plant recognition (Jermy, 1966). These deterrent or “bitter” compounds are usually plant secondary metabolites, which have diverse molecular structures. Such compounds inhibit food intake in the majority of phytophagous insects, except for some specialized species that use them as token stimuli (Schoonhoven et al., 2005).

Phytophagous insects have sophisticated taste systems to recognize deterrent compounds, which direct their feeding behavior (Yarmolinsky et al., 2009). The sense of taste is located in sensilla that are mainly situated on the mouthparts, tarsi, ovipositor, and antennae (Bernays and Chapman, 1994). These sensilla take the form of hairs or cones with a terminal pore in the cuticular structure, and often contain the dendrites of three or four gustatory sensory neurons (GSNs). These GSNs are directly connected to the central nervous system (CNS) with no intervening synapses. Analyses using the tip-recording technique for taste sensilla led to the discovery of a “deterrent neuron” in the larvae of *Bombyx mori* (Ishikawa, 1966) and *Pieris brassicae* (Ma, 1969). Since then, GSNs coding for feeding deterrents have been identified in maxillary sensilla in larvae and tarsal sensilla of adults of many Lepidopteran species. However, the molecular basis of these deterrent neurons remains unclear.

Gustatory receptors (GRs) expressed in the dendrites of GSNs determine the selectivity of the response of GSNs (Thorne et al., 2004; Wang et al., 2004). Since the first insect GRs were identified in the model organism *Drosophila melanogaster* (Clyne et al., 2000), the function of some of its bitter GRs have been revealed (Dweck and Carlson, 2020; Freeman and Dahanukar, 2015). Five *D. melanogaster* GRs (Gr47a, Gr32a, Gr33a, Gr66a, and Gr22e) are involved in sensing strychnine (Lee et al., 2010; Lee et al., 2015; Moon et al., 2009; Poudel et al., 2017). Nicotine-induced action potentials are dependent on Gr10a, Gr32a, and Gr33a (Rimal and Lee, 2019). Gr8a, Gr66a, and Gr98b function together in the detection of L-canavanine (Shim et al., 2015). Gr33a, Gr66a, and Gr93a participate in the responses to caffeine and umbelliferone (Lee et al., 2009; Moon et al., 2006; Poudel et al., 2015). Gr28b is necessary for avoiding saponin (Sang et al., 2019). However, only a few studies have focused on the function of bitter GRs in phytophagous insects. PxutGr1, a bitter GR in *Papilio xuthus*, was found to respond specifically to the oviposition stimulant, synephrine (Ozaki et al., 2011). Most recently, Gr66, a bitter GR in *B. mori,* was reported to be responsible for the mulberry-specific feeding preference of silkworms. To date, no studies have validated the functions of GRs coded to respond to feeding and oviposition deterrents in phytophagous insects (Zhang et al., 2019).

*Plutella xylostella* (L.) is the most widespread Lepidopteran pest species, causing losses of US$ 4–5 billion per year (You et al., 2020; Zalucki et al., 2012). It has developed resistance to the usual insecticides because of its short life cycle (14 days) (Furlong et al., 2013). *P. xylostella* mainly selects *Brassica* species as its host plants. Aliphatic or indole glucosinolates as well as their hydrolyzed products (e.g., isothiocyanates or nitriles) are oviposition attractants and stimulants for females (Renwick et al. 2006, Sun et al. 2009). Brassinolide (BL) is a C_28_ brassinosteroid (BR) that exhibits the highest activity among all BRs tested to date (Clouse and Sasse, 1998; Grove et al., 1979; Mitchell et al., 1970). BRs occur ubiquitously in the plant kingdom, and are present in almost every plant part including leaves and pollen. Exogenous application of BL and its analog 24-epibrassinolide (EBL) to plants has been shown to increase their stress resistance (Clouse and Sasse, 1998). Interestingly, BL has a strikingly similar structure to that of ecdysteroid hormones in arthropods, such as 20-hydroxyecdysone (Fujioka and Sakurai, 1997). Their structures are so similar that BL shows agonistic activity with 20-hydroxyecdysone in many insect species (Zullo and Adam, 2002; Smagghe et al., 2002). However, it is still unclear whether BL can be perceived by the gustatory system of insects.

In this study, we demonstrate that *P. xylostella* can sense the plant hormone BL as a deterrent. We identified one bitter GR (PxylGr34) by analyzing the *P. xylostella* genome (Engsontia et al., 2014; You et al., 2013), our unpublished data, and transcriptome data from Yang et al. (2017). Our results show that PxylGr34 is highly expressed in the larval head and the female antennae, and is strongly tuned to the plant hormone BL, which suppresses larval feeding and ovipositing of mated females of *P. xylostella*.

## Results

### Identification of the GR gene *PxylGr34* in *P. xylostella*

By analyzing the transcriptome data obtained from the antennae, forelegs (only tibia and tarsi), and head (without antennae) of adults, and the mouthparts of 4^th^ instar larvae, we rechecked the 79 GR sequences reported in previous genomic (Engsontia et al., 2014; You et al., 2013) and transcriptomic studies (Yang et al., 2017) of *P. xylostella*. After removing repetitive sequences and deleting paired sequences with amino acid identity greater than 99%, we obtained 67 GR sequences (Figure 1—figure supplement 1 and Supplementary file 1). Next, we calculated the transcripts per million (TPM) values of these GRs (Figure 1—figure supplement 2). The TPM value of *PxylGr34*, which clustered with bitter GRs, was much higher than those of other GR genes in the antennae, head and forelegs, as well as the larval mouthparts (Figure 1—figure supplement 2). *PxylGr34* was originally identified from genomic data (Engsontia et al., 2014), and then detected in the antennal transcriptomic data of *P. xylostella* (named *PxylGr2* in Yang et al., 2017). However, both studies provided only its partial coding sequences (Figure 1—figure supplement 3). Based on genomic data and our transcriptomic data, we obtained the full-length coding sequence of *PxylGr34* through gene cloning and Sanger sequencing (Figure 1—figure supplement 3). The protein encoded by *PxylGr34* is a typical characteristic GR with seven transmembrane domains, and a full open reading frame (ORF) of 418 amino acids (Figure 1—figure supplement 4).

### High PxylGr34 transcript levels in the larval head and female antenna

To explore the expression patterns of *PxylGr34* in the larvae and adults, we detected its relative transcript levels in different tissues of 4^th^ instar larvae and females using quantitative real-time PCR (qPCR). *P. xylostella* is most damaging to host plants at the 4^th^ instar stage (Harcourt, 1957). We detected high *PxylGr34* transcript levels in the larval head. We also detected *PxylGr34* transcripts in the larval thorax and gut (Figure 1A). In the adult females, *PxylGr34* transcripts were restricted to the antennae (Figure 1B).

**Figure 1.**
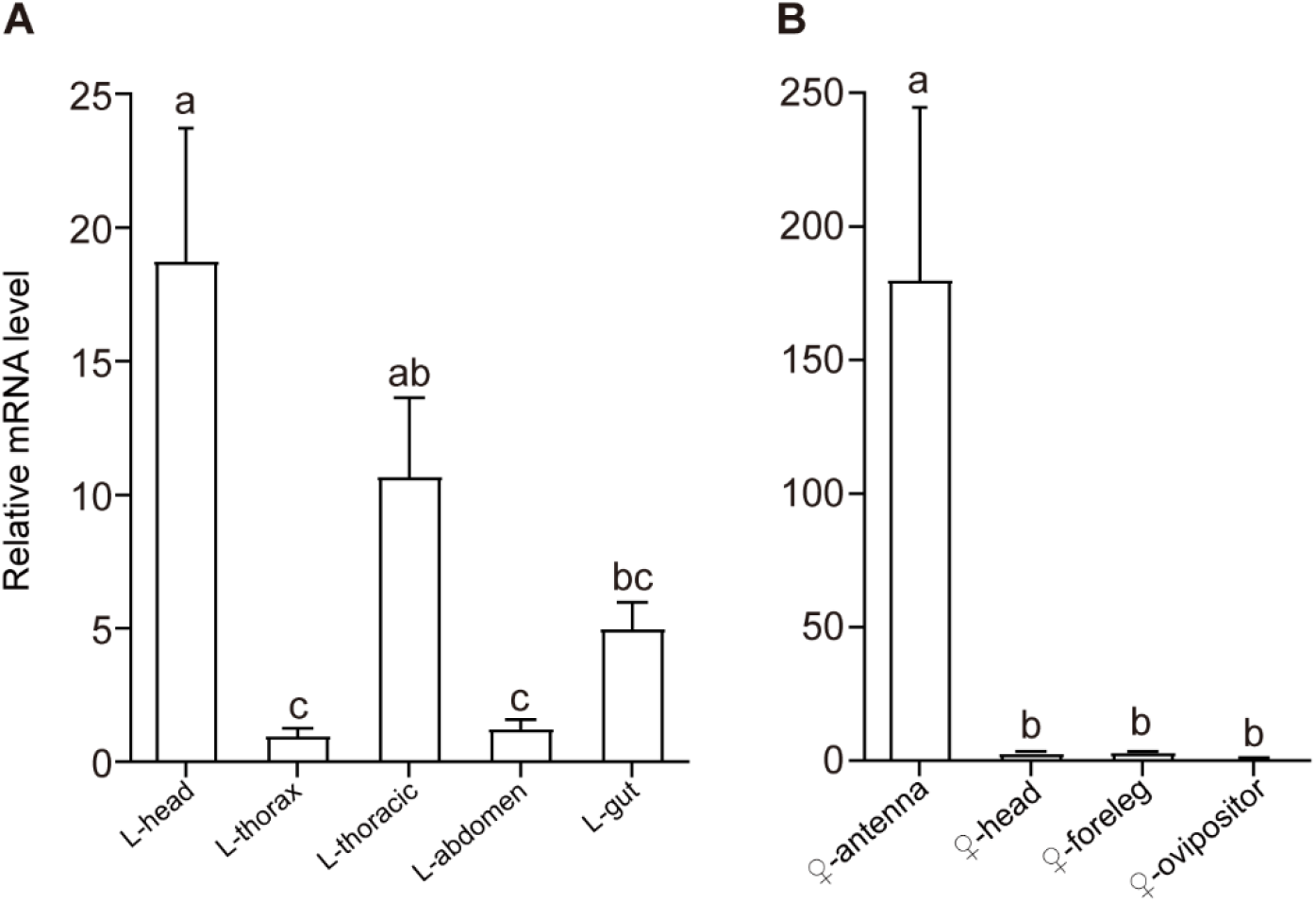
High PxylGr34 transcript levels in the larval head and female antenna. (**A**) Relative *PxylGr34* transcript levels in different tissues of 4^th^ instar larvae of *Plutella xylostella* as determined by qPCR: L-head, larval head; L-thorax, larval thorax (without wing, thoracic, gut, or other internal tissues); L-thoracic, larval thoracic; L-abdomen, larval abdomen (without gut or other internal tissues); L-gut, larval gut. (**B**) Relative *PxylGr34* transcript levels in different tissues of 4^th^ instar larvae of *Plutella xylostella* as determined by qPCR: ♀-antenna, female antenna; ♀-head, female head (without antennae); ♀-foreleg, female foreleg (only tarsi and tibia); ♀-ovipositor, female ovipositor. Data are mean ± SEM. *n* = 3 replicates of 40–200 tissues each. For 4^th^ instar larvae, *F*_(4, 10)_ = 14.1, *p* = 0.0004; for mated females, *F*_(3, 8)_ = 24.44, *p* = 0.0002 (one-way ANOVA, Tukey’s HSD test).

### BL and its analog EBL induced a strong response in the oocytes expressing PxylGr34

We used the *Xenopus laevis* oocyte expression system and two-electrode voltage-clamp recording to study the function of PxylGr34. Among 24 tested phytochemicals including sugars, amines, amino acids, plant hormones, and secondary metabolites, BL induced a strong response in the oocytes expressing PxylGr34, as did its analog EBL at a concentration of 10^−4^ M (Figure 2A and Figure 2B). The currents induced by BL increased from the lowest threshold concentration of 3.3×10^−5^ M in a dose-dependent manner (Figure 2C). Oocytes expressing PxylGr34 showed weak responses to methyl jasmonate and allyl isothiocyanate, but no response to 20-hydroxyecdysone and other tested compounds (Figure 2A and Figure 2B). As negative controls, the water-injected oocytes failed to respond to any of the tested chemical stimuli (Figure 2—figure supplement 1).

**Figure 2.**
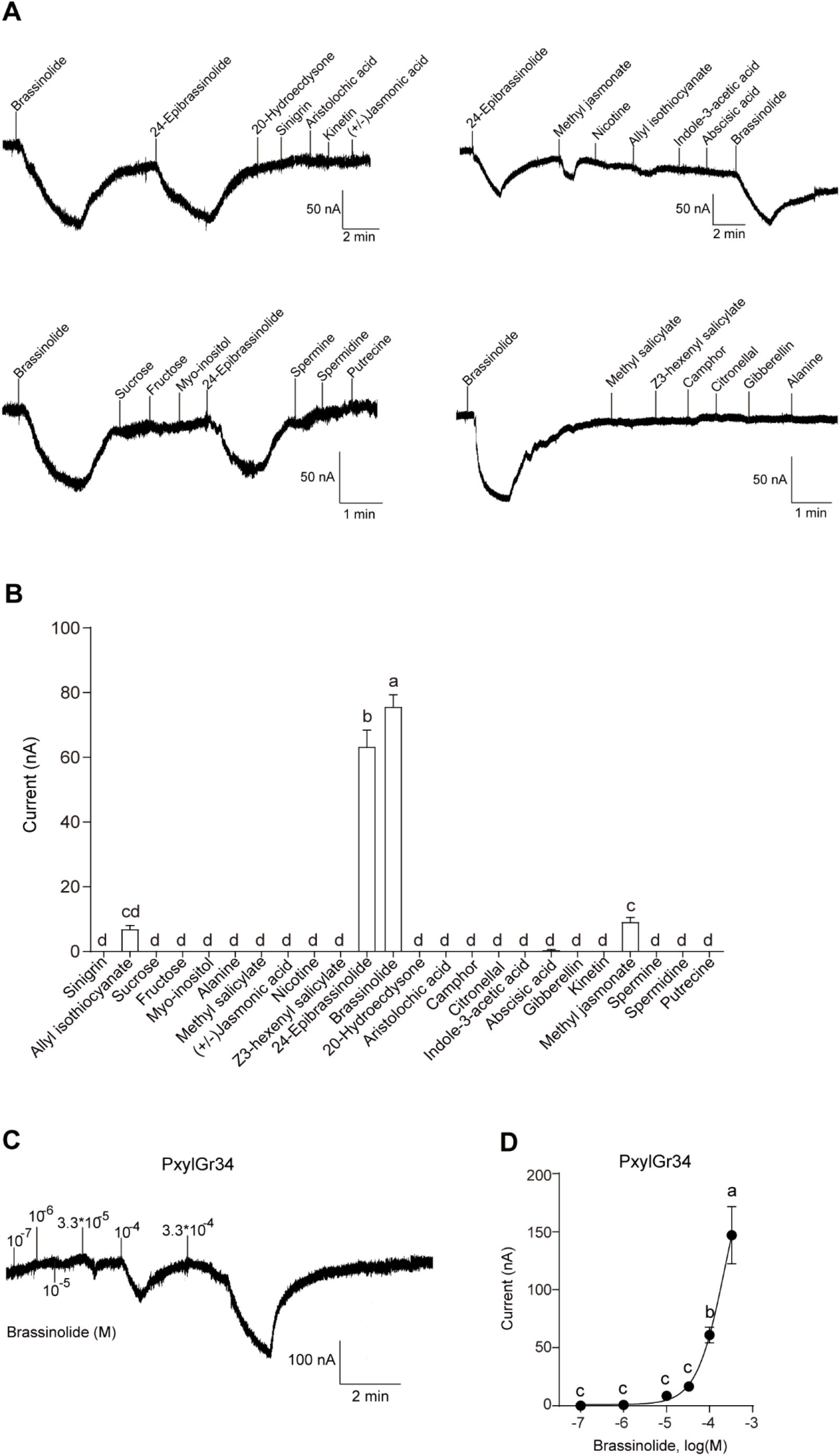
BL and its analog EBL induced a strong response in the oocytes expressing PxylGr34. (**A**) Inward current responses of *Xenopus* oocytes expressing PxylGr34 in response to ligands at 10^−4^ M. (**B**) Response profiles of *Xenopus* oocytes expressing PxylGr34 in response to ligands at 10^−4^ M. Data are mean ± SEM. *n* = 7 replicates of cells. *F*_(23, 144)_ = 202.8, *p* < 0.0001 (one-way ANOVA, Tukey HSD test). (**C**) Inward current responses of *Xenopus* oocytes expressing PxylGr34 in response to BL at a range of concentrations. (**D**) Response profiles of *Xenopus* oocytes expressing PxylGr34 in response to BL at a range of concentrations. Data are mean ± SEM; *n* = 6–8 replicates of cells. *F*_(5, 38)_ = 31.36, *p* < 0.0001 (one-way ANOVA, Tukey’s HSD test).

### The larval medial sensilla styloconica exhibited vigorous responses in *P. xylostella* to BL

Next, using the tip recording technique, we examined whether any gustatory sensilla in the mouthparts of larvae of *P. xylostella* could respond to BL. Of the two pairs of sensilla styloconica in the 4^th^ instar larvae, the medial sensilla styloconica exhibited vigorous responses to BL at 3.3×10^−4^ M, while the lateral sensilla styloconica had no response (Figure 3A and Figure 3B). The medial sensilla styloconica showed a dose-dependent response to BL, although the testing concentrations were limited because of the low solubility of BL in water (highest concentration approximately 3.3×10^−4^ M) (Figure 3C and Figure 3D).

**Figure 3.**
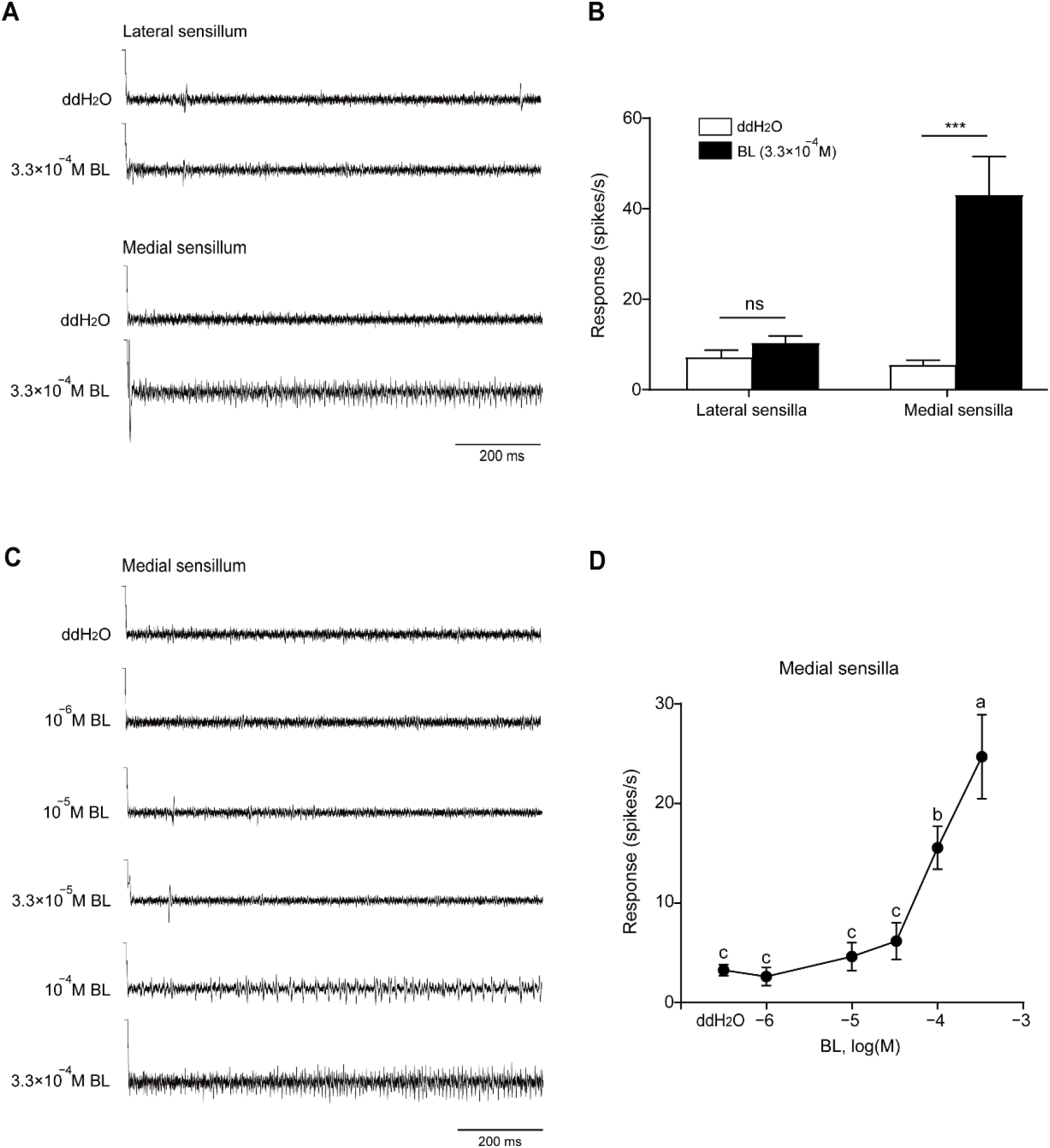
The larval medial sensilla styloconica exhibited vigorous responses in *P. xylostella* to BL. (**A**) Typical electrophysiological recordings in response to water and BL at 3.3×10^−4^ M for 1 s obtained by tip-recording from a neuron innervating the lateral and medial sensillum styloconica on maxillary galea of *P. xylostella* 4^th^ instar larvae. (**B**) Response profiles of the two lateral and medial sensilla styloconica on the maxilla of *P. xylostella* fourth instar larvae to water and BL at 3.3×10^−4^ M. For lateral sensilla, *n* = 14 replicates of larvae, *t*_(13)_ = 1.282, *p* = 0.2223; for medial sensilla, *n* = 16 replicates of larvae, *t*_(15)_ = 4.763, *p* = 0.0003. Data are mean ± SEM. Data were analyzed by paired-samples *t*-test. (**C**) Typical electrophysiological recordings in response to water and BL at a series of concentrations for 1 s obtained by tip-recording from a neuron innervating the medial sensillum styloconicum on the maxillary galea of *P. xylostella* 4^th^ instar larvae. (**D**) Response profiles of medial sensilla styloconica on maxilla of *P. xylostella* 4^th^ instar larvae to water and BL at a series of concentrations. Data are mean ± SEM. *n* = 8–20 replicates of larvae, *F*_(5, 84)_ =17.58, *p* < 0.0001 (one-way ANOVA, Tukey’s HSD test).

### BL and EBL induced feeding deterrence effect on *P. xylostella* larvae

We further tested the effects of BL on the larval feeding behavior of *P. xylostella*. In a dual-choice feeding test with 4^th^ instar larvae, the feeding areas of larvae were significantly smaller on the leaf discs treated with BL at concentrations of 10^−4^ M and above than on the control leaf discs. In addition, the feeding preference index of larvae to BL decreased with increasing BL concentrations (Figure 4A). Similar results were obtained with EBL (Figure 4B), indicating that both BL and EBL function as feeding deterrents to *P. xylostella* larvae. The survival rate of the larvae after 24 h in all the treatments in the test was greater than 95%, indicating that BL and EBL do not have direct toxic effects on larvae.

**Figure 4.**
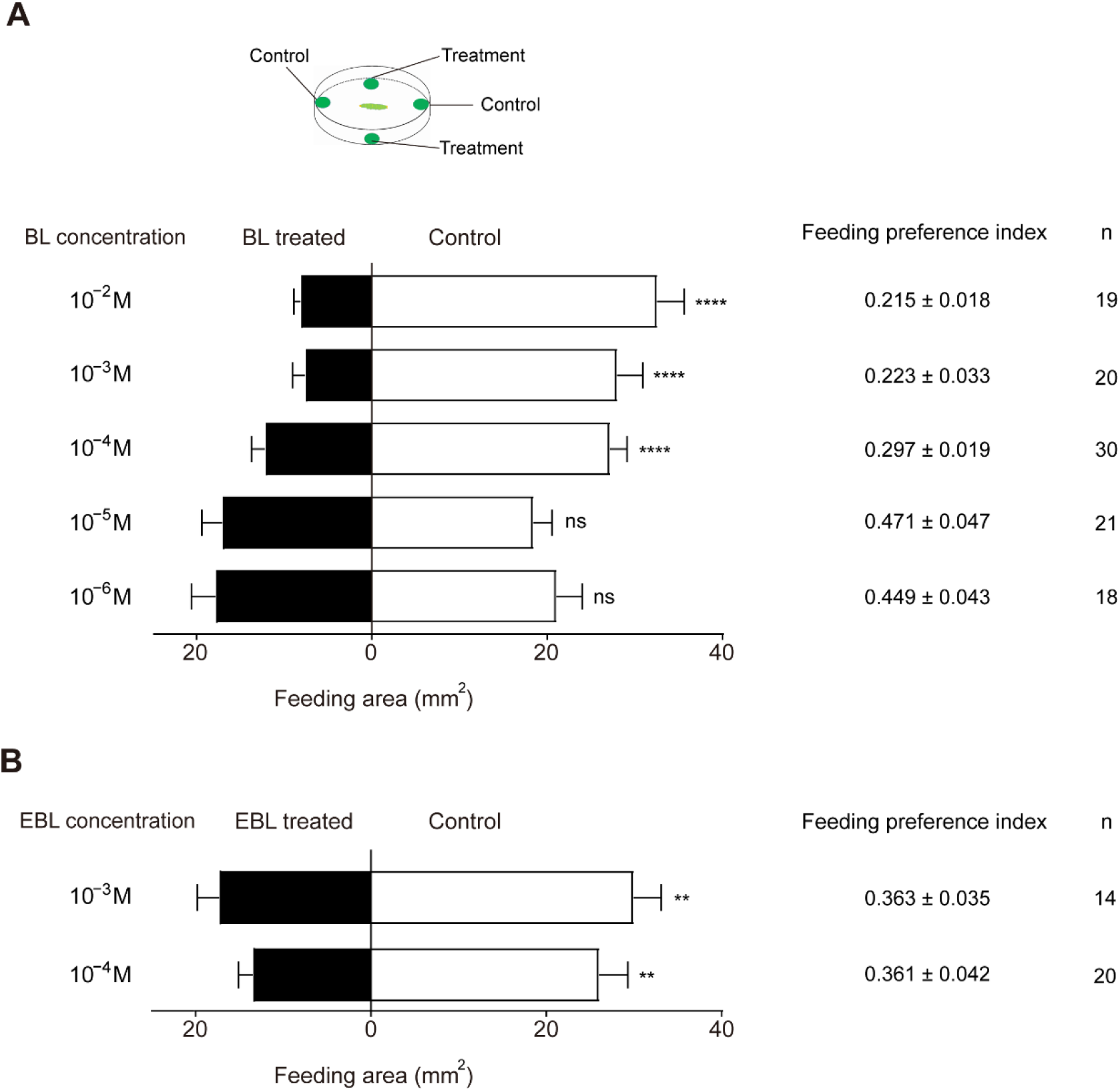
BL and EBL Induced Feeding Deterrence Effect on *P. xylostella* Larvae. Dual-choice feeding tests were conducted using pea leaf discs. (**A**) Total feeding area of control (white bars) and BL-treated discs (black bars). BL was diluted in 50% ethanol to 10^−6^, 10^−5^, 10^−4^, 10^−3^, and 10^−2^ M. For 10^−6^ M, *n* = 18 replicates of larvae, *t*_(17)_ =0.8733, *p* = 0.3947; for 10^−5^ M, *n* = 21 replicates of larvae, *t*_(20)_ = 0.5378, *p* = 0.5967; for 10^−4^ M, *n* = 30 replicates of larvae, *t*_(29)_ = 8.582, *p* < 0.0001; for 10^−3^ M, *n* = 20 replicates of larvae, *t*_(19)_ = 6.012, *p* < 0.0001; for 10^−2^ M, *n* = 19 replicates of larvae, *t*_(18)_ = 7.873, *p* < 0.0001. Data are mean ± SEM. Data were analyzed by paired-samples *t*-test. Preference index (mean ± SEM) and simple linear regression were calculated based on feeding areas (see Methods for details). (**B**) Total feeding area of control (white bars) and EBL-treated discs (black bars). EBL was diluted in 50% ethanol at 10^−4^ and 10^−3^ M. For 10^−4^ M, *n* = 20 replicates of larvae, *t*_(19)_ = 3.497, *p* = 0.0024; 10^−3^ M, *n* = 14 replicates of larvae, *t*_(13)_ = 3.778, *p* = 0.0023. Data are mean ± SEM. Data were analyzed by paired-samples *t*-test.

### PxylGr34 siRNA treated *P. xylostella* larvae alleviated the feeding deterrent effect of BL

To clarify whether PxylGr34 mediates the behavioral responses of *P. xylostella* larvae to BL *in vivo*, we tested the effect of siRNA targeting *PxylGr34* on the feeding behavior of the 4^th^ instar larvae. The relative transcript level of *PxylGr34* in the head of larvae treated with PxylGr34 siRNA was half that in the head of larvae treated with green fluorescent protein (GFP) siRNA or ddH_2_O (Figure 5A). This confirmed that feeding with siRNA is an effective method for RNAi-inhibition of *PxylGr34* in the larval head. To test the effects of RNAi-knockdown of *PxylGr34* on the feeding behavior of 4^th^ instar larvae, the siRNA-treated larvae were subjected to a dual-choice leaf disc feeding assay as described above, with leaf discs of pea (*Pisum sativum* L.) treated with 10^−4^ M BL or untreated (control). As shown in Figure 5B, both the water-treated larvae and the GFP siRNA-treated larvae preferred control leaf discs over those treated with BL, while the RNAi-treated larvae showed no significant preference (Figure 5B). Thus, the knock-down of PxylGr34 by RNAi alleviated the deterrent effect of BL on the feeding of *P. xylostella* larvae.

**Figure 5.**
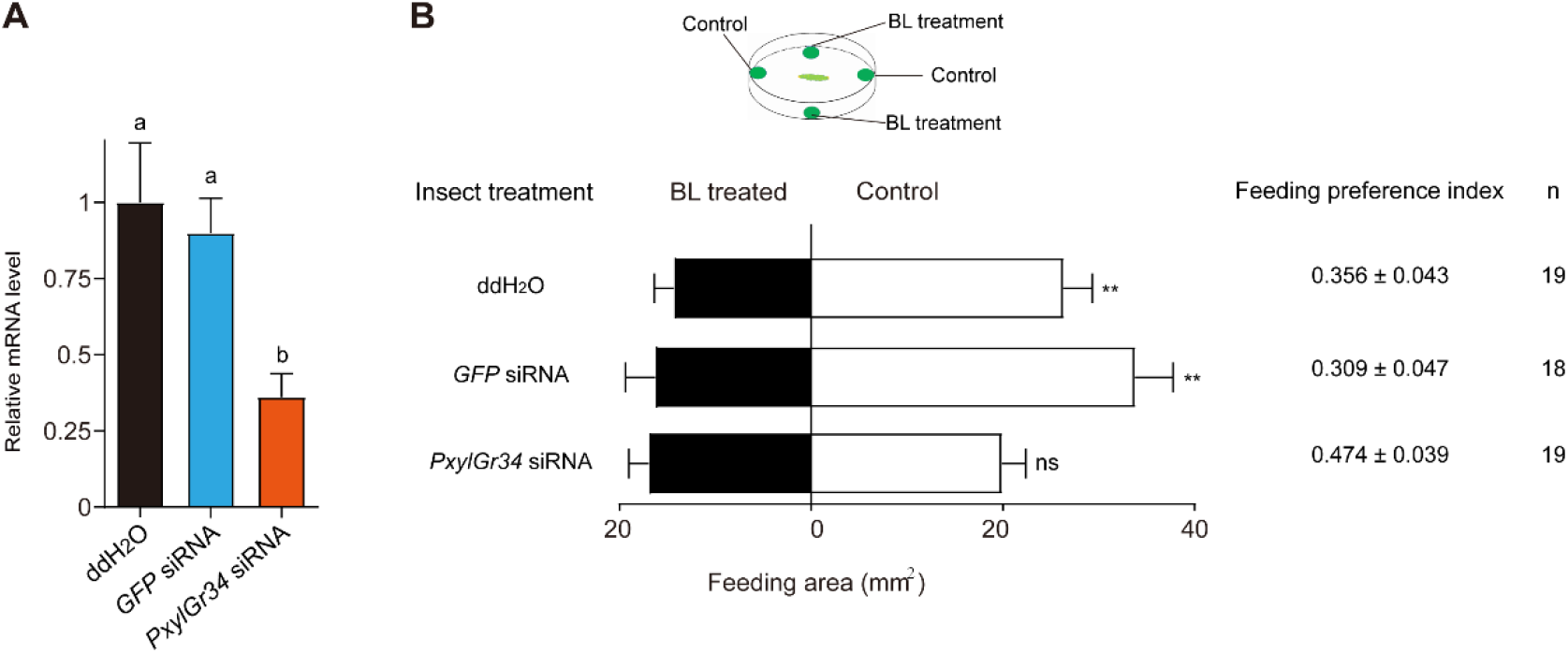
PxylGr34 siRNA treated *P. xylostella* larvae alleviated the feeding deterrent effect of BL. (**A**) Effect of PxylGr34 siRNA on transcript levels of *PxylGr34* in 4^th^ instar larval head. Heads were collected after feeding larvae with cabbage leaf discs coated with *PxylGr34* siRNA, *GFP* siRNA, or ddH_2_O. Relative transcript levels of *PxylGr34* in each treatment were determined by qPCR. *n* = 3 replicates of 21–24 heads each. Data are mean ± SEM. *F*_(2, 6)_ = 7.443, *p* = 0.0237 (one-way ANOVA, Tukey’s HSD test). (**B**) Choice assay using 4^th^ instar larvae fed on cabbage leaf discs treated with *PxylGr34* siRNA, *GFP* siRNA, or ddH_2_O. In dual choice assay, larvae chose between control pea leaf discs (treated with 50% ethanol) and those treated with BL at 10^−4^ M (diluted in 50% ethanol). Figure shows total feeding area of control (white bars) and treated discs (black bars). For ddH_2_O treatment, *n* = 19 replicates of larvae, *t*_(18)_ = 2.912, *p* = 0.0093; for *GFP* siRNA treatment, *n* = 18 replicates of larvae, *t*_(17)_ =3.28, *p* = 0.0044; *PxylGr34* siRNA treatment, *n* = 19 replicates of larvae, *t*_(18)_ =1.374, *p* = 0.1864. Data are mean ± SEM. Data were analyzed by paired-samples *t*-test. Feeding preference index (mean ± SEM) was calculated based on feeding area (see Methods for details).

### BL induced oviposition deterrence to *P. xylostella* females

We also tested the effects of BL on the female ovipositing behavior of *P. xylostella*. In a dual-choice oviposition test with mated females, significantly fewer eggs were laid on the sites treated with BL at 10^−4^ M, 10^−3^ M, and 10^−2^ M than on the control sites. In addition, the oviposition preference index decreased as the BL concentrations increased (Figure 6).

**Figure 6.**
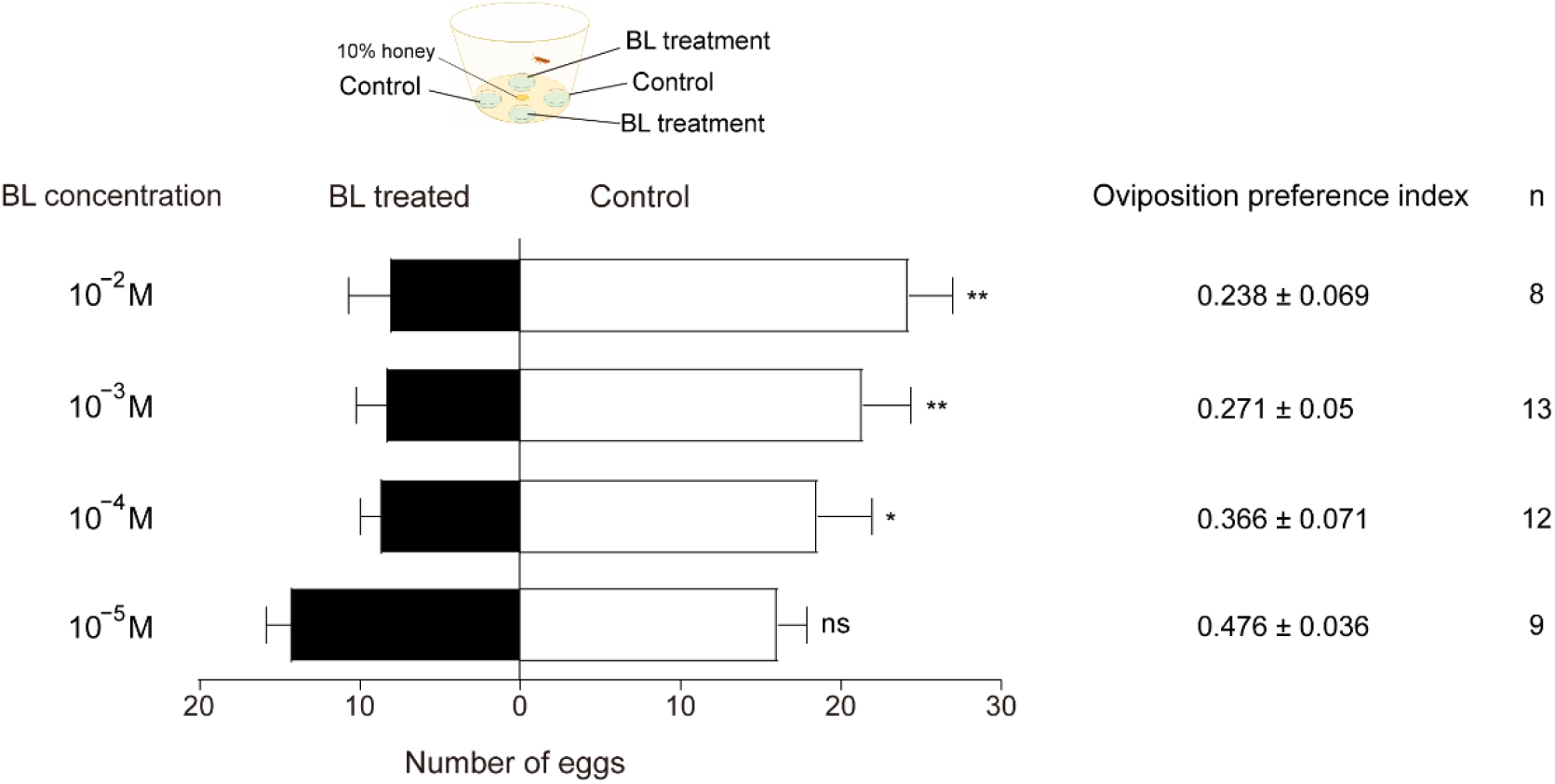
BL induced oviposition deterrence to *P. xylostella* females. Dual-choice oviposition tests were conducted using plastic film coated with cabbage leaf juice. Both control (50% ethanol) and BL (diluted in 50% ethanol) were painted evenly onto plastic film. After 24 h, total number of eggs laid by a single mated female on control films (white bars) and BL-treated films (black bars) were counted. For 10^−5^ M, *n* = 9 replicates of female moths, *t*_(8)_ = 0.8192, *p* = 0.4364; for 10^−4^ M, *n* = 12 replicates of female moths, *t*_(11)_ = 2.539, *p* = 0.0275; for 10^−3^ M, *n* = 13 replicates of female moths, *t*_(12)_ = 4.072, *p* = 0.0015; 10^−2^ M, *n* = 8 replicates of female moths, *t*_(7)_ = 4.249, *p* = 0.0038. Data are mean ± SEM. Data were analyzed using paired-samples *t*-test. Oviposition preference index (mean ± SEM) and simple linear regressions were calculated from number of eggs laid (see Materials and Methods for details).

## Discussion

In this study, we identified the full-length coding sequence of *PxylGr34* from our transcriptome data, and found that this gene is highly expressed in the head of the 4^th^ instar larvae and in the antennae of females. Our results show that PxylGr34 is specifically tuned to BL and its analog EBL, and that the medial sensilla styloconica of 4^th^ instar larvae have electrophysiological responses to BL. Our results also show that BL inhibits larval feeding and female ovipositing of *P. xylostella*, and that knock-down of PxylGr34 by RNAi can abolish the feeding inhibition effect of BL. This is the first study to show that an insect can detect and react to this steroid plant hormone. The results of the systematic functional analyses of the GR, the electrophysiological responses of the sensilla, behavior, and behavioral regulation *in vivo* show that PxylGr34 is a bitter GR specifically tuned to BL. This receptor mediates the deterrent effects of BL on feeding and ovipositing behaviors of *P. xylostella*.

The larvae of Lepidopteran species have two pairs of gustatory sensilla (medial and lateral sensilla styloconica) located in the maxillae galea; these sensilla play a decisive role in larval food selection (Dethier, 1937; Schoonhoven and van Loon, 2002). Each sensillum usually contains four gustatory sensory neurons, of which one is often responsive to deterrents. Ishikawa (1966) described a “deterrent neuron” in the medial sensillum styloconicum of silkworm, *Bombyx mori*, and showed that it responds to several plant alkaloids and phenolics (Ishikawa, 1966). Similar neurons have been found in other Lepidopteran species, but their profiles vary. For example, the tobacco hornworm, *Manduca sexta*, has a deterrent neuron in the medial sensillum styloconicum that responds to aristolochic acid, and another deterrent neuron in the lateral sensillum styloconicum that responds to salicin, caffeine, and aristolochic acid (Glendinning et al., 2002). The diversity of deterrent neurons facilitates the detection of, and discrimination among, a wide range of diverse taste stimuli, implying that different bitter GRs are expressed in these neurons. In a previous study, one GSN in the medial maxillary sensillum styloconicum of the 4^th^ instar larvae of the diamondback moth was found to respond to sinigrin and glucosinolates (van Loon et al., 2002). In this study, we identified one neuron responding to BL in the same sensillum. For *P. xylostella* larvae, sinigrin is a feeding stimulant while BL is a feeding deterrent. We speculate that the GRs tuned to sinigrin and BL are located in different GRNs in the medial sensilla styloconica.

The GR family is massively expanded in moth species, and most of the GRs are bitter GRs (Cheng et al., 2017). However, few studies have functionally characterized bitter GRs. In *D. melanogaster*, the loss of bitter GRs was found to eliminate repellent behavior in response to specific noxious compounds. For example, Gr33a mutant flies could not avoid non-volatile repellents like quinine and caffeine (Moon et al., 2009), and mutation of Gr98b impaired the detection of L-canavanine (Shim et al., 2015). When the bitter GR PxutGr1 of *P. xuthus* was knocked-down by RNAi, the oviposition behavior in response to synephrine was strongly reduced (Ozaki et al., 2011). Knock-out of the bitter GR BmorGr66 in *B. mori* larvae resulted in a loss of feeding specificity for mulberry (Zhang et al., 2019). In this study, knock-down of PxylGr34 in the larvae of *P. xylostella* eliminated the feeding deterrence of BL. These results indicate that this bitter GR is specifically tuned to BL and mediates the aversive response of larvae to BL and related compounds.

Brassinolide was first isolated from rape pollen in *Brassica napus* L. in 1979 (Grove et al., 1979), and it was the first brassinosteroid (BR) hormone to be discovered in plants. Since then, many BRs have been found throughout the plant kingdom. In plants, BRs promote elongation and stimulate cell division, participate in vascular differentiation and fertilization, and affect senescence (Clouse and Sasse, 1998). Several studies have shown that treatment with exogenous BRs can enhance plants’ tolerance to chilling stress, high temperatures, and salt, and improve resistance to various pathogens (Clouse and Sasse, 1998). The first BL receptor identified in plants was brassinosteroid insensitive 1 (BRI1) in *Arabidopsis thaliana* (Li and Chory, 1997). As a leucine-rich repeat receptor-like kinase (LRR-RLK), BRI1 is localized on the plasma membrane (Friedrichsen et al. 2000), is ubiquitously expressed in different plant organs, and shares homology with other receptor kinases in plants and animals (Li and Chory, 1997). In the term of its response selectivity, BRI1 is less sensitive to other BRs such as castasterone than to BL, and is insensitive to 20-hydroxyecdysone (Wang et al., 2001).

As the first reported BL receptor in animals, PxylGr34 is specifically tuned to BL and EBL, similar to the selectivity of the BL receptor BRI1 in plants. However, compared with BRI1, PxylGr34 is much less sensitive to BL (Kd value ~10^8^ M) (Wang et al., 2001). PxylGr34 with its putative seven-transmembrane domain (Figure 1—figure supplement 4) belongs to the group of bitter GRs that may function as ligand-gated ion channels (Sato et al., 2011). Therefore, the BL receptors in plants and animals do not have similar structures, but their functional convergence reflects the importance of BL in the co-evolution of plants and phytophagous insects.

Both BL and EBL show striking structural similarities to the ecdysteroid-type arthropod hormone, 20-hydroxyecdysone (20E). However, they cannot replace the function of 20E to induce insect ecdysis (Richter and Koolman, 1991). However, in one study, injection with 20 μg 24-EBL was fatal to mid last-instar larvae of *Spodoptera littoralis* (Smagghe et al., 2002). The BL content differs widely among plant species; for example, it is 1.37×10^−4^ g/kg in *Brassica campestris* L. leaves and 1.25×10^−6^ g/kg in *Arabidopsis thaliana* leaves (Lv et al., 2014). Our results show that the threshold concentration of BL for behavioral inhibition of *P. xylostella* is in the range of 10^−4^–10^−3^ g/kg, suggesting that BL functions as a plant hormone at low concentrations, but as an insect deterrent at higher concentrations.

There is a rich variety of bitter GRs in phytophagous insects, but only a few have been functionally characterized (Kasubuchi et al., 2018; Zhang et al., 2019). In this study, we showed that PxylGr34, a bitter GR highly expressed in larval head and female antennae of *P. xylostella*, is tuned to the plant hormones BL and EBL, and that this interaction mediates the aversive feeding/oviposition responses of *P. xylostella* to these compounds. These findings not only increase our understanding of the gustatory coding mechanisms of feeding/oviposition deterrents in phytophagous insects, but also offer new perspectives for using plant hormones as potential agents to suppress pest insects.

## Materials and Methods

### Insects and plants

*P. xylostella* was obtained from the Institute of Plant Protection, Shanxi Academy of Agricultural Sciences, China. The insects were reared at the Institute of Zoology, Chinese Academy of Sciences, Beijing. The larvae were fed with cabbage (*Brassica oleracea* L.) and kept at 26 ± 1°C with a 16L:8D photoperiod and 55% –65% relative humidity. The diet for adults was a 10% (v/v) honey solution. Pea (*Pisum sativum* L.) plants were grown in an artificial climate chamber at 26 ± 1°C with a 16L:8D photoperiod and 55%–65% relative humidity. The plants were grown in nutrient soil in pots (8 × 8 × 10 cm), and were 4–5 weeks old when they were used in experiments.

### Care and use of *Xenopus laevis*

All procedures were approved by the Animal Care and Use Committee of the Institute of Zoology, Chinese Academy of Sciences, and followed The Guidelines for the Care and Use of Laboratory Animals (protocol number IOZ17090-A). Female *X. laevis* were provided by Prof. Qing-Hua Tao (MOE Key Laboratory of Protein Sciences, Tsinghua University, China) and reared in our laboratory with pig liver as food. Six healthy naive *X. laevis* 18–24 months of age were used in these experiments. They were group-housed in a box with purified water at 20 ± 1°C. Before experiments, each *X. laevis* individual was anesthetized by bathing in an ice–water mixture for 30 min before surgically collecting the oocytes.

### Sequencing and expression analysis of GR genes in *P. xylostella*

We conducted transcriptome analyses of the *P. xylostella* moth antennae, foreleg (only tibia and tarsi), head (without antenna), and the 4^th^ instar larval mouthparts. Total RNA was extracted using QIAzol Lysis Reagent (Qiagen, Hilden, Germany) and treated with DNase I following the manufacturer’s protocol. Poly(A) mRNA was isolated using oligo dT beads. First-strand complementary DNA was generated using random hexamer-primed reverse transcription, followed by synthesis of second-strand cDNA using RNaseH and DNA polymerase I. Paired-end RNA-seq libraries were prepared following Illumina’s protocols and sequenced on the Illumina HiSeq 2500 platform (Illumina, San Diego, CA). High-quality clean reads were obtained by removing adaptors and low-quality reads, then *de novo* assembled using the software package Trinity v2.8.5 (Haas et al., 2013). The GR genes were annotated by NCBI BLASTX searches against a pooled insect GR database, including the previously reported *P. xylostella* GRs (Engsontia et al., 2014; You et al., 2013; Yang et al., 2017). The translated amino acid sequences of the identified GRs were further aligned manually by NCBI BLASTP and tools at the T-Coffee web server (http://tcoffee.crg.cat/apps/tcoffee/do:expresso). The TPM values were calculated using the software package RSEM v1.2.28 (Li and Dewey, 2011) to analyze GR gene transcript levels.

### Phylogenetic analysis

Phylogenetic analysis of *P. xylostella* GRs was performed based on amino acid sequences, together with those of previously reported GRs of *Heliconius Melpomene* (Briscoe et al., 2013) and *B. mori* (Guo et al., 2017). The phylogenetic tree was constructed using MEGA6.0 (RRID: SCR_000667) with the neighbor-joining method and the p-distances model (Tamura et al., 2013).

### RNA isolation and cDNA synthesis

The tissues were dissected, immediately placed in a 1.5-mL Eppendorf™ tube containing liquid nitrogen, and stored at −80°C until use. Total RNA was extracted using QIAzol Lysis Reagent following the manufacturer’s protocol (including DNase I treatment), and RNA quality was checked with a spectrophotometer (NanoDrop™ 2000; Thermo Fisher Scientific, Waltham, MA, USA). The single-stranded cDNA templates were synthesized using 2 μg total RNAs from various samples with 1 μg oligo (dT) 15 primer (Promega, Madison, WI, USA). The mixture was heated to 70°C for 5 min to melt the secondary structure of the template, then M-MLV reverse transcriptase (Promega) was added and the mixture was incubated at 42°C for 1 h. The products were stored at −20°C until use.

### Cloning of *PxylGr34* from *P. xylostella*

Based on the candidate full-length nucleotide sequences of *PxylGr34* identified from our transcriptome data, we designed specific primers (Supplementary file 1). All amplification reactions were performed using Q5 High-Fidelity DNA Polymerase (New England Biolabs, Beverly, MA, USA). The PCR conditions for amplification of *PxylGr34* were as follows: 98°C for 30 s, followed by 30 cycles of 98°C for 10 s, 60°C for 30 s, and 72°C for 1 min, and final extension at 72° C for 2 min. Templates were obtained from antennae of female *P. xylostella*. The sequences were further verified by Sanger sequencing.

### Quantitative real-time PCR

The qPCR analyses were conducted using the QuantStudio 3 Real-Time PCR System (Thermo Fisher Scientific) with SYBR Premix Ex *Taq*™ (TaKaRa, Shiga, Japan). The gene-specific primers to amplify an 80–150 bp product were designed by Primer-BLAST (http://www.ncbi.nlm.nih.gov/tools/primer-blast/) (Supplementary file 1). The thermal cycling conditions were as follows: 10 s at 95°C, followed by 40 cycles of 95°C for 5 s and 60°C for 31 s, followed by a melting curve analysis (55°C – 95°C) to detect a single gene-specific peak and confirm the absence of primer dimers. The product was verified by nucleotide sequencing. *PxylActin* (GenBank number: AB282645.1) was used as the control gene (Teng et al., 2012). Each reaction was run in triplicate (technical replicates) and the means and standard errors were obtained from three biological replicates. The relative copy numbers of *PxylGr34* were calculated using the 2^−ΔΔCt^ method (Livak and Schmittgen, 2001).

### Receptor functional analysis

The full-length coding sequence of PxylGr34 was first cloned into the pGEM-T vector (Promega) and then subcloned into the pCS2^+^ vector. cRNA was synthesized from the linearized modified pCS2^+^ vector with mMESSAGE mMACHINE SP6 (Ambion, Austin, TX, USA). Mature healthy oocytes were treated with 2 mg mL^−1^ collagenase type I (Sigma-Aldrich, St Louis, MO, USA) in Ca^2+^-free saline solution (82.5 mM NaCl, 2 mM KCl, 1 mM MgCl_2_, 5 mM HEPES, pH = 7.5) for 20 min at room temperature. Oocytes were later microinjected with 55.2 ng cRNA. Distilled water was microinjected into oocytes as the negative control. Injected oocytes were incubated for 3–5 days at 16°C in a bath solution (96 mM NaCl, 2 mM KCl, 1 mM MgCl_2_, 1.8 mM CaCl_2_, 5 mM HEPES, pH = 7.5) supplemented with 100 mg mL^−1^ gentamycin and 550 mg mL^−1^ sodium pyruvate. Whole-cell currents were recorded with a two-electrode voltage clamp. The intracellular glass electrodes were filled with 3 M KCl and had resistances of 0.2–2.0 MΩ. Signals were amplified with an OC-725C amplifier (Warner Instruments, Hamden, CT, USA) at a holding potential of −80 mV, low-pass filtered at 50 Hz, and digitized at 1 kHz. Data were acquired and analyzed using Digidata 1322A and pCLAMP software (RRID: SCR_011323) (Axon Instruments Inc., Foster City, CA, USA). Dose-response data were analyzed using GraphPad Prism software (RRID: SCR_002798 6) (GraphPad Software Inc., San Diego, CA, USA).

### Electrophysiological responses of contact chemosensilla on the maxilla of larvae to BL

The tip-recording technique was used to record action potentials from the lateral and medial sensilla styloconica on the maxillary galea of larvae, following the protocols described elsewhere (Hodgson et al., 1955; van Loon, 1990; van Loon et al., 2002). Distilled water served as the control stimulus, as KCl solutions elicited considerable responses from the galeal styloconic taste sensilla (van Loon et al., 2002).

Experiments were carried out with larvae that were 1–2 days into their final stadium (4^th^ instar). The larvae were starved for 15 min before analysis. The larvae were cut at the mesothorax, and then silver wire was placed in contact with the insect tissue. The wire was connected to a preamplifier with a copper miniconnector. A glass capillary filled with the test compound, into which a silver wire was inserted, was placed in contact with the sensilla. Electrophysiological responses were quantified by counting the number of spikes in the first second after the start of stimulation. The interval between two successive stimulations was at least 3 min to avoid adaptation of the tested sensilla. Before each stimulation, a piece of filter paper was used to absorb the solution from the tip of the glass capillary containing the stimulus solution to avoid an increase in concentration due to evaporation of water from the capillary tip. The temperature during recording ranged from 22° to 25°C. Action potentials (spikes) were amplified by an amplifier (Syntech Taste Probe DTP-1, Hilversum, The Netherlands) and filtered (A/D-interface, Syntech IDAC-4). The electrophysiological signals were recorded by SAPID Tools software version 16.0 (Smith et al., 1990), and analyzed using Autospike software version 3.7 (Syntech).

### Dual-choice feeding bioassays

Dual-choice feeding assays were used to quantify the behavioral responses of *P. xylostella* larvae to BL and EBL. These assays were based on the protocol reported by van Loon et al. (2002), with modifications. The axisymmetric pinnate leaf was freshly picked from 4-week-old pea plants grown in a climate-controlled room. One leaf was folded in half, and two leaf discs (diameter, 7 mm) were punched from the two halves as the control (C) and treated (T) discs, respectively. For the treated discs, 5 μL (13 μL/cm^2^) of the test compound diluted in 50% ethanol was spread on the upper surface using a paint brush. For the control discs, 5 μL 50% ethanol was applied in the same way. Control (C) and treated (T) discs were placed in a C-T-C-T sequence around the circumference of the culture dish (60 mm diameter × 15 mm depth; Corning, NY, USA). After the ethanol had evaporated, a single 4^th^ instar caterpillar (day 1), which had been starved for 6 h, was placed in each dish. The dishes were kept for 24 h at 23°–25°C in the dark, to avoid visual stimuli. Each dish was covered with a circular filter paper disc (diameter 7 cm) moistened with 200 μL ddH_2_O to maintain humidity. At the end of the test, the leaf discs were scanned using a DR-F120 scanner (Canon, Tokyo, Japan) and the remaining leaf area was quantified with ImageJ software (NIH). Paired-samples *t*-test was used to detect differences in the consumed leaf area between control and treated leaf discs. The feeding preference index (FPI) values were calculated according to the following equation: (consumed area of treated disc)/ (consumed area of treated disc + consumed area of control disc). FPI values of 1 and 0 indicate complete preference or deterrence for the test compound, respectively, and an FPI of 0.5 indicates no preference.

### Dual-choice oviposition bioassays

A dual-choice oviposition bioassay was used to quantify the behavioral responses of *P. xylostella* mated female moths to BL. This assay was modified from the protocol reported by Gupta and Thorsteinson (1960) and Justus and Mitchell (1996). A paper cup (10 cm diameter × 8 cm height) with a transparent plastic lid (with 36 pinholes for ventilation) was used for ovipositing of the mated females. Fresh cabbage leaf juice was centrifuged at 3000 rpm for 5 min, and 60 μL of the supernatant was spread with a paintbrush onto PE film from clinical gloves. Four culture dishes (35 mm diameter) (Corning, New York, NY, USA) covered with these PE films were placed on the bottom of each cup. This oviposition system was developed based on the biology of *P. xylostella* (Harcourt, 1957). On each of two diagonally positioned treatment films, 125 μL BL (13 μL/ cm^2^) diluted in 50% ethanol was evenly spread on the upper surface using a paint brush. On the other two diagonally positioned films, 125 μL 50% ethanol (control) was spread in the same way. After the ethanol had evaporated, a small piece of absorbent cotton soaked with 10% honey-water mixture was placed in the center of the cup.

The pupae of *P. xylostella* were selected and newly emerged adults were checked and placed in a mesh cage (25 × 25 × 25 cm), with a 10% honey-water mixture supplied during the light phase. The female:male ratio was 1:3 to ensure that all the females would be mated. After at least 24 h of mating time, the mated females were removed from the cage and placed individually into the oviposition cup during the light phase. After 24 h at 26 ± 1°C with a 16L: 8D photoperiod and 55%–65% relative humidity, the number of eggs on each plastic film was counted. Paired-samples *t*-test was used to detect significant differences in the number of eggs laid between the control and treatment films. The oviposition preference index (OPI) values were calculated as follows: OPI = (no. of eggs on treated film)/ (no. of eggs on treatment film + no. of eggs on control film). OPI values of 1 and 0 indicate complete preference and complete deterrence for the test compound, respectively, and an OPI of 0.5 indicates no preference.

### siRNA preparation

A unique siRNA region specific to *PxylGr34* was selected guided by the siRNA Design Methods and Protocols (Humana Press, 2013). The siRNA was prepared using the T7 RiboMAX™ Express RNAi System kit (Promega, Madison) following manufacturer’s protocol. The GFP (GenBank: M62653.1) siRNA was designed and synthesized using the same methods. We tested three different siRNAs of *PxylGr34* based on different sequence regions, and selected the most effective and stable one for further analyses. The oligonucleotides used to prepare siRNAs are listed in Supplementary file 2.

### Oral delivery of siRNA

The siRNAs were supplied to the larvae by oral delivery as reported elsewhere (Chaitanya et al., 2017; Gong et al., 2011), with some modifications. Each siRNA was spread onto cabbage leaf discs (*Brassica oleracea*) and fed to 4^th^-instar larvae. Freshly punched cabbage leaves discs (diameter 0.7 cm) were placed into 24-well clear multiple well plates (Corning, NY, USA). For each disc, 3 μg siRNA diluted in 6 μL 50% ethanol was evenly distributed on the upper surface using a paint brush. Both 50% ethanol and GFP siRNA were used as negative controls. After the ethanol had evaporated, one freshly molted 4^th^-instar larva, which had been starved for 6 h, was carefully transferred onto each disc and then allowed to feed for 12 h. Each well was covered with dry tissue paper to maintain humidity. The larvae that had consumed the entire disc were selected and starved for another 6 h, and then these larvae were used in the dual choice behavioral assay or for qPCR analyses as described above. The larvae that did not consume the treated discs were discarded.

### Data analysis

Data were analyzed using GraphPad Prism 8.3. Figures were created using GraphPad Prism 8.3 and Adobe illustrator CS6 (RRID: SCR_014198) (Adobe systems, San Jose, CA). Two-electrode voltage-clamp recordings, electrophysiological dose-response curves, and the square-root transformed qPCR data were analyzed by one-way ANOVA and Tukey’s HSD tests with two distribution tails. These analyses were performed using GraphPad prism 8.3. Electrophysiological response data and all dual-choice test data were analyzed using the two-tailed paired-samples *t*-test. Simple linear regressions were calculated by GraphPad prism 8.3. Statistical tests and the numbers of replicates are provided in the figure legends. In all statistical analyses, differences were considered significant at *p* < 0.05. Asterisks represent significance: * *p* < 0.05, ** *p* < 0.01, *** *p* < 0.001, **** *p* < 0.0001; ns, not significant. Response values are indicated as means ± SEM; and *n* represents the number of samples in all cases.

## Acknowledgements

We thank the lab members Hao Guo, Nan-Ji Jiang, Rui Tang, and Jun Yang for their helps in the data analysis and comments, Shuai-Shuai Zhang, Yan Chen and Ruo-Xi Shi for their assistances in tip-recording analysis. We thank Xi-Zhong Yan from Shanxi Agricultural University for the assistance in insect rearing, Prof. Qing-Hua Tao from MOE Key Laboratory of Protein Sciences, Tsinghua University for providing *Xenopus laevis* frogs. We also thank Prof. Bill Hansson from Max Planck Institute for Chemical Ecology, Germany, for his comments on this work. This work is funded by the National Natural Science Foundation of China (Grant No. 31830088), National Key R&D Program of China (Grant No. 2017YFD0200400), and China Postdoctoral Science Foundation (Grant No. 2019M660792).

---

**Figure 1—figure supplement 1.**
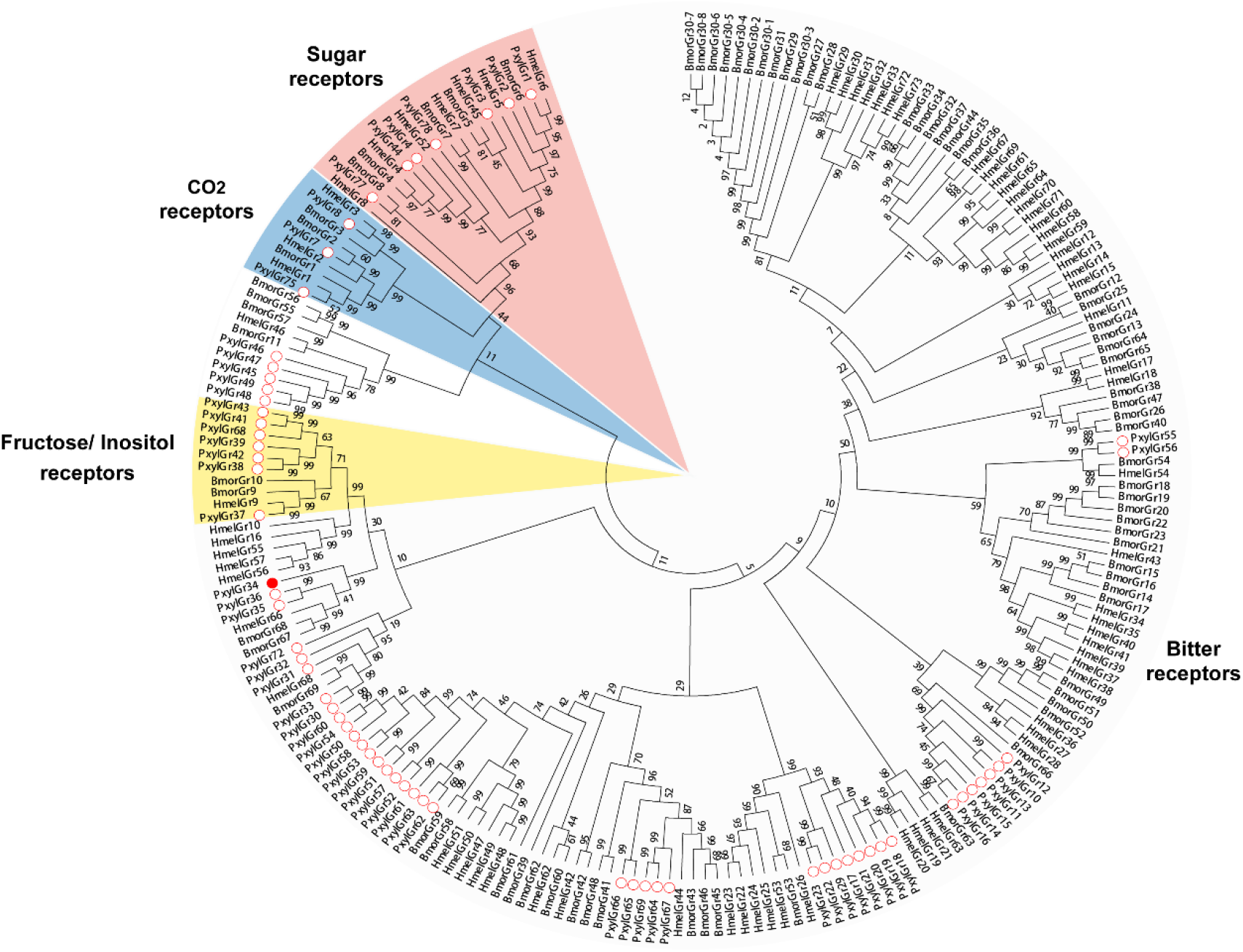
Phylogenetic tree of gustatory receptors (GRs). Amino acid sequences are based on previously reported transcriptome data for functionally identified GRs. Bootstrap values are based on 1,000 replicates. Bar indicates phylogenetic distance. Abbreviations: Hmel, *Heliconius melpomene*; Bmor, *Bombyx mori*; Pxyl, *Plutella xylostella*. ○, GRs of *P. xylostella*; ●, PxylGR34.

**Figure 1—figure supplement 2.**
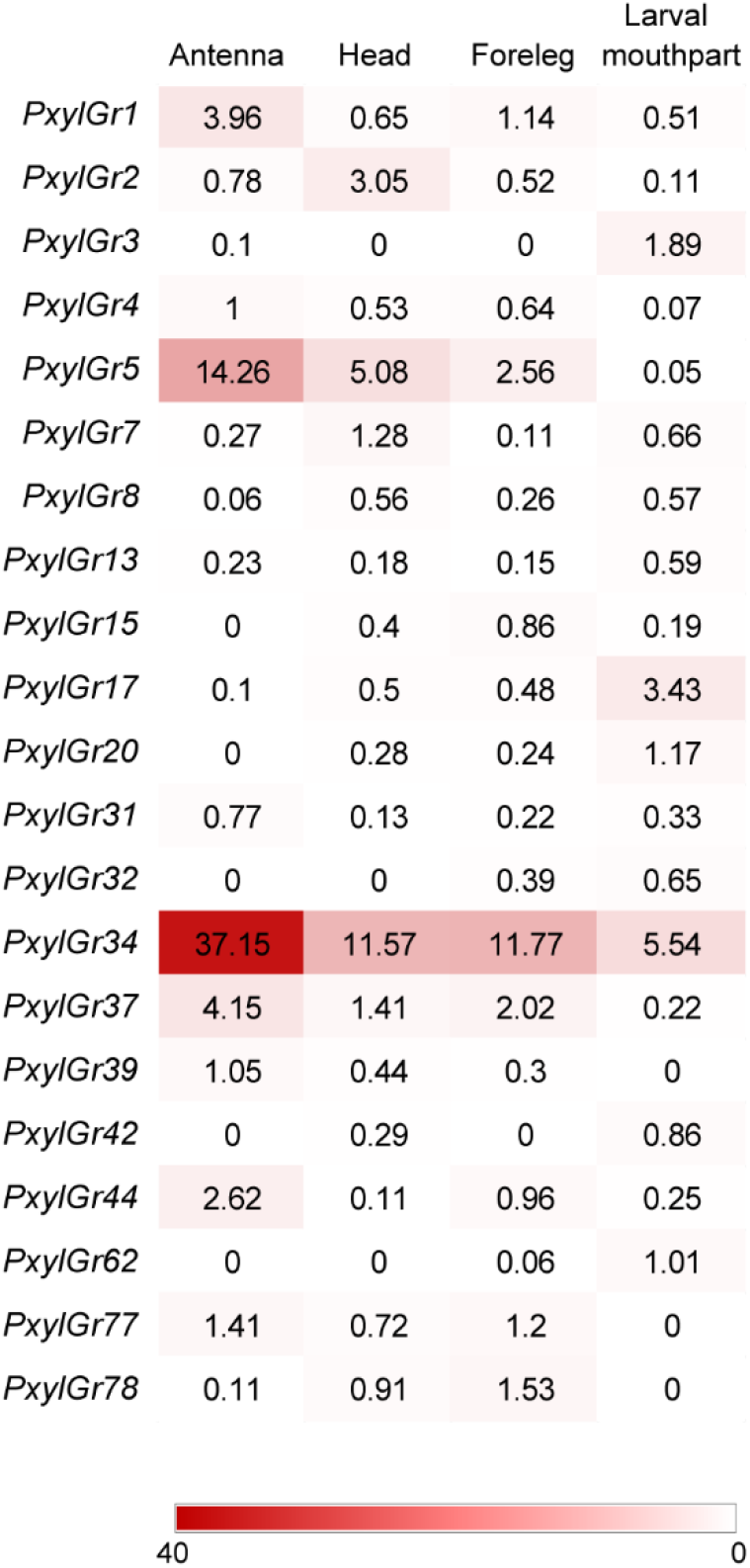
Tissue expression pattern of gustatory receptors (GRs) in *Plutella xylostella* as determined by Illumina read-mapping analysis. Transcripts per million (TPM) value of each GR is indicated in box. Color scales were generated using Microsoft Excel. Antenna, head, and foreleg were from the moth; larval mouthpart was from 4^th^ instar larvae.

**Figure 1—figure supplement 3.**
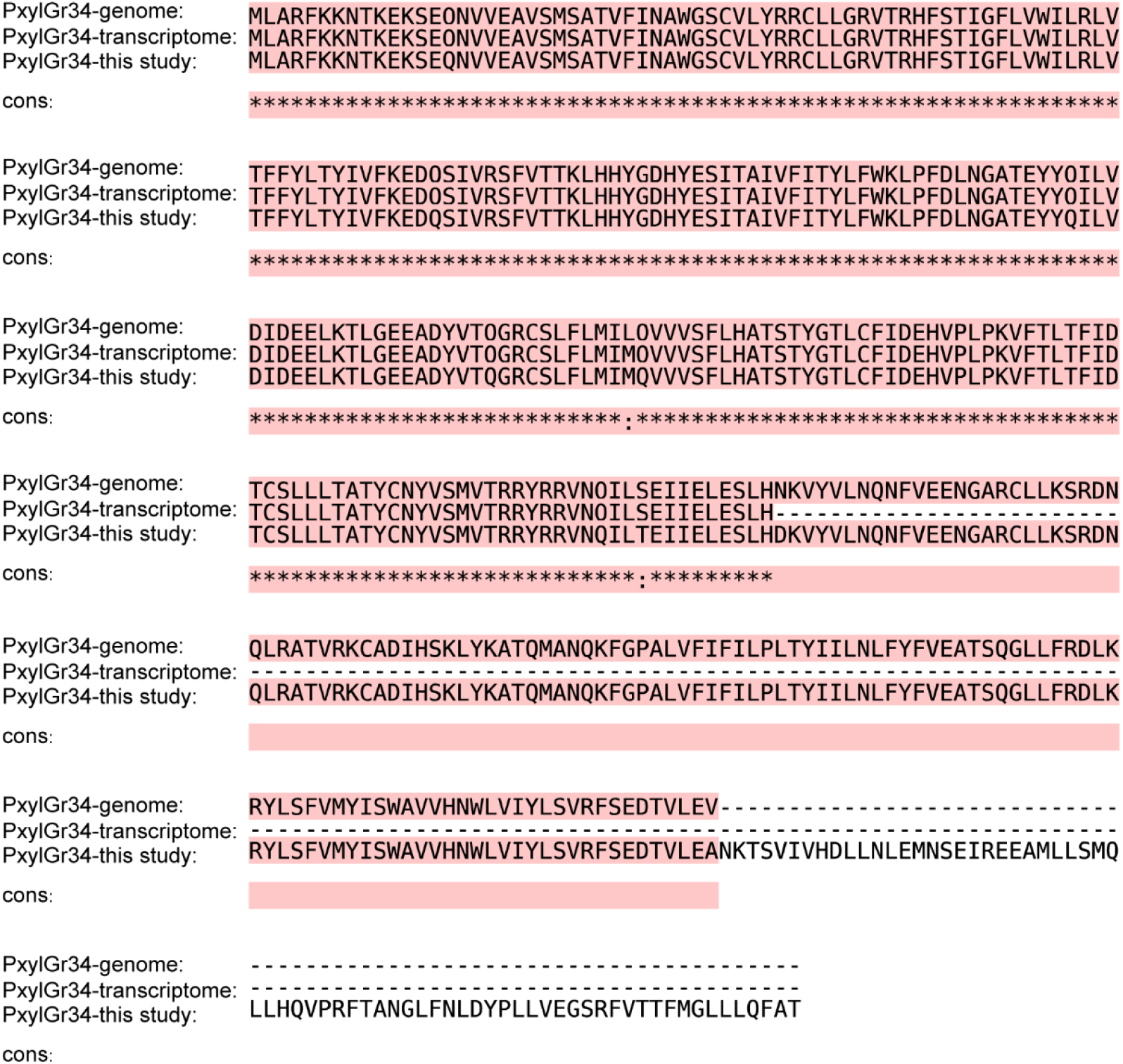
Alignment of amino acid sequences of PxylGr34 from genomic data (Engsontia et al., 2014), transcriptomic data (Yang et al., 2017), and this study.

**Figure 1—figure supplement 4.**
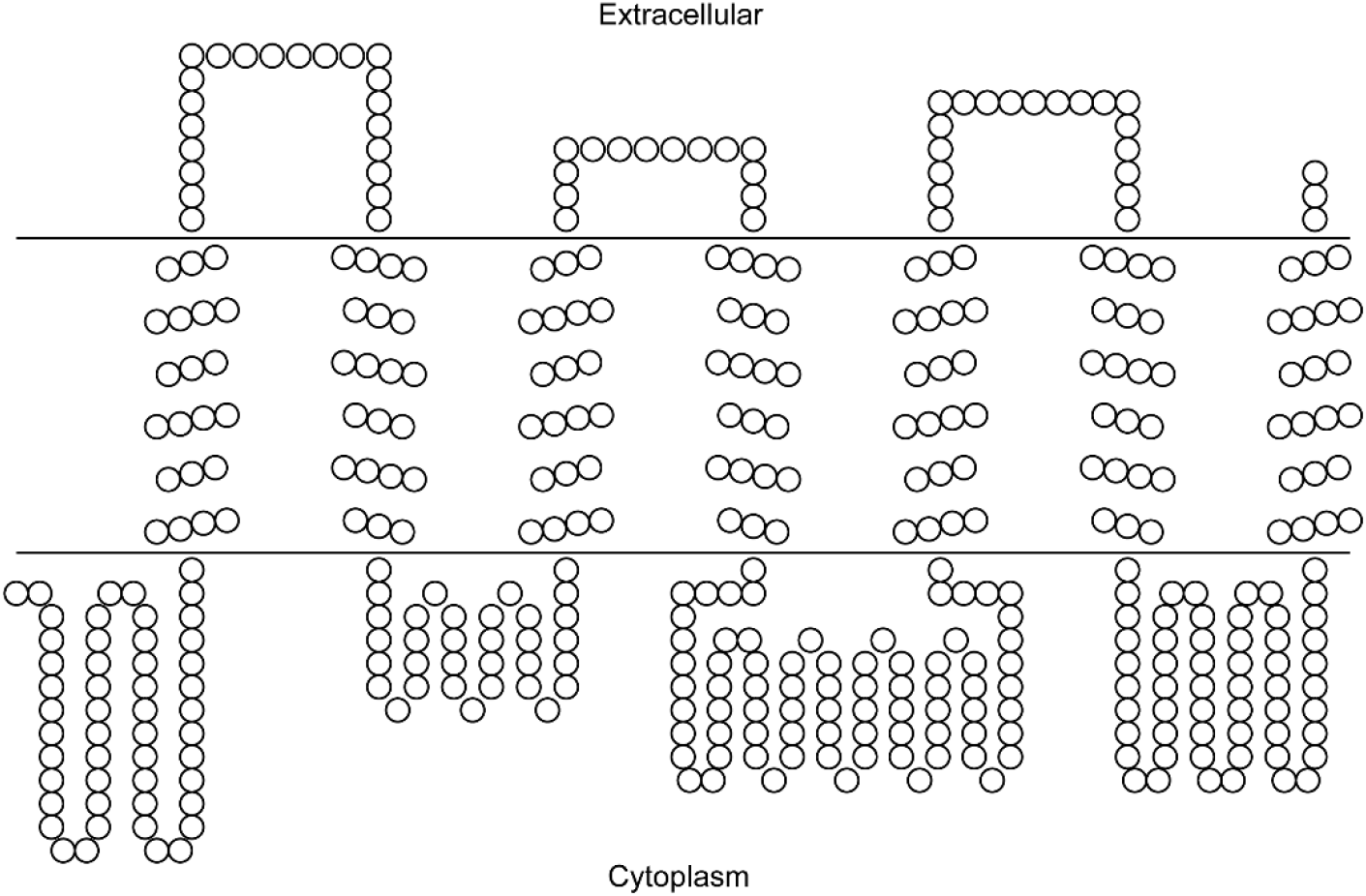
Secondary structure prediction of PxylGr34. Image was constructed using TOPO2 software (http://www.sacs.ucsf.edu/TOPO2/) based on secondary structure predicted by TOPCONS (topcons.net) models (Tsirigos et al., 2015). Only the model with a reliable seven-transmembrane structure was adopted.

**Figure 2—figure supplement 1.**
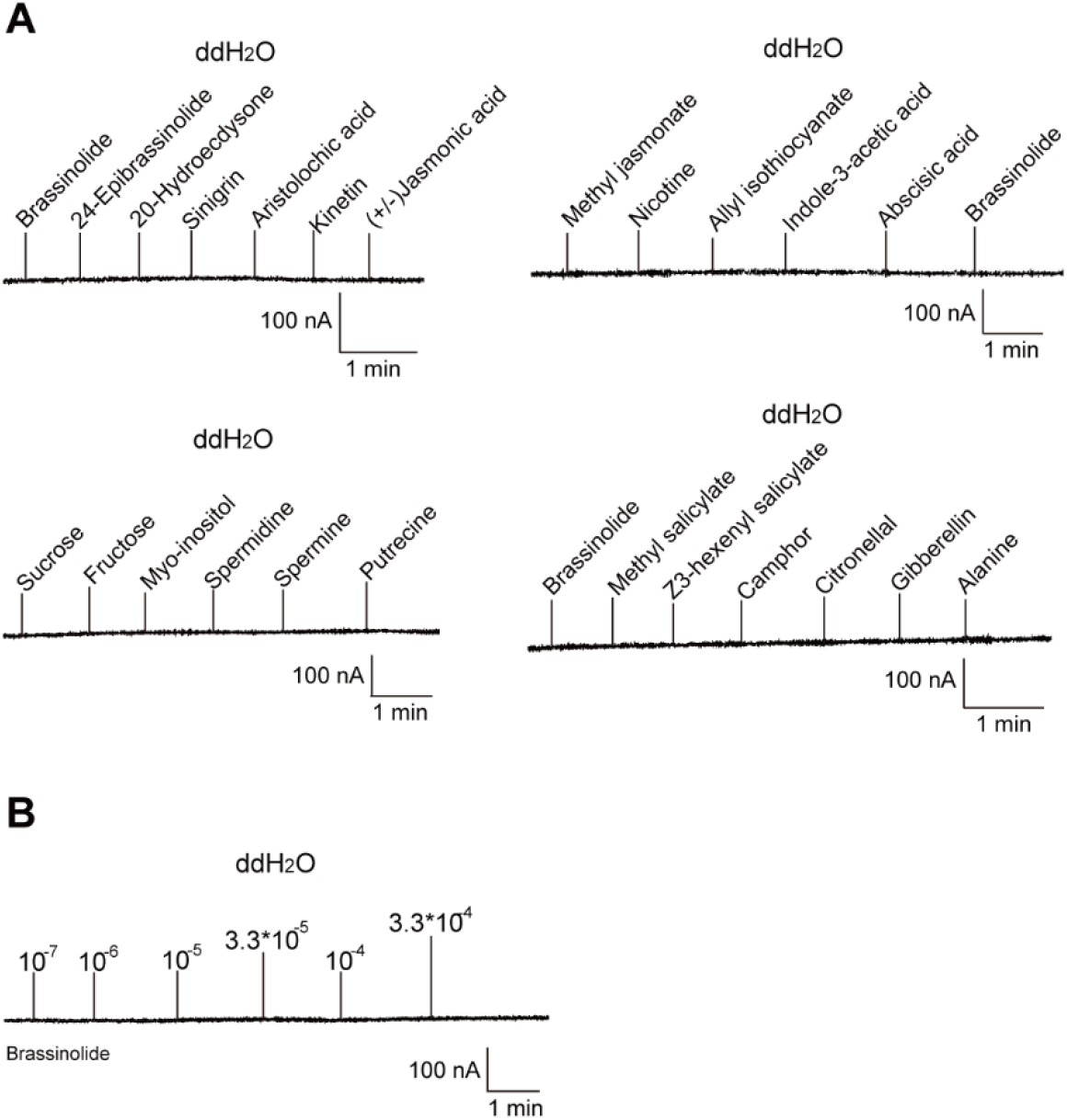
Two-electrode voltage-clamp recordings of *Xenopus* oocytes injected with distilled water and stimulated with test compounds. No inward current responses of distilled-water-injected *Xenopus* oocytes to tested ligands (**A**) or BL at a range of concentrations (**B**).

## Supplementary files

**Supplementary file 1.**Sequence information for gustatory receptors of *Plutella xylostella*.

**Supplementary file 2.**Primers used for qPCR, *Xenopus* oocyte expression (Xe), and siRNA synthesis.

**Table S1.** Sequence Information for Gustatory Receptors of *Plutella xylostella*.

| Reported PxylGrs | PxylGrs in this study | GenBank number | Notes |
| --- | --- | --- | --- |
| PxylGr1* | PxylGr1 | XP_011560061.1 |  |
| PxylGr2* | PxylGr2 | XP_011560059.1 |  |
| PxylGr3* | PxylGr3 | XP_011560063.1 |  |
| PxylGr4* | PxylGr4 | XP_011560065.1 |  |
| PxylGr5* | (delete) | XP_011568607.1 | Partial sequence of PxylGr75 |
| PxylGr6* | (delete) |  | Partial sequence of PxylGr75 |
| PxylGr7* | PxylGr7 | XP_011557481.1 |  |
| PxylGr8* | PxylGr8 | XP_011558222.1 |  |
| PxylGr9* | (delete) |  | Partial sequence of PxylGr8 |
| PxylGr10* | PxylGr10 |  |  |
| PxylGr11* | PxylGr11 |  |  |
| PxylGr12* | PxylGr12 |  |  |
| PxylGr13* | PxylGr13 |  |  |
| PxylGr14* | PxylGr14 |  |  |
| PxylGr15* | PxylGr15 |  |  |
| PxylGr16* | PxylGr16 |  |  |
| PxylGr17* | PxylGr17 |  |  |
| PxylGr18* | PxylGr18 |  |  |
| PxylGr19* | PxylGr19 |  |  |
| PxylGr20* | PxylGr20 |  |  |
| PxylGr21* | PxylGr21 |  |  |
| PxylGr22* | PxylGr22 |  |  |
| PxylGr23* | PxylGr23 |  |  |
| PxylGr24* | (delete) |  | Partial sequence of PxylGr19 |
| PxylGr25* | (delete) |  | Repetitive sequence with PxylGr20 |
| PxylGr26* | (delete) |  | Partial sequence of PxylGr21 |
| PxylGr27* | (delete) |  | Repetitive sequence with PxylGr22 |
| PxylGr28* | (delete) |  | Repetitive sequence with PxylGr23 |
| PxylGr29* | PxylGr29 |  |  |
| PxylGr30* | PxylGr30 | XP_011561237.1 |  |
| PxylGr31* | PxylGr31 | XP_011561224.1 |  |
| PxylGr32* | PxylGr32 |  |  |
| PxylGr33* | PxylGr33 |  |  |
| PxylGr34* | PxylGr34 |  |  |
| PxylGr35* | PxylGr35 | XP_011565113.1 |  |
| PxylGr36* | PxylGr36 |  |  |
| PxylGr37* | PxylGr37 | XP_011548483.1 |  |
| PxylGr38* | PxylGr38 | XP_011552106.1 |  |
| PxylGr39* | PxylGr39 |  |  |
| PxylGr40* | PxylGr40 |  |  |
| PxylGr41* | PxylGr41 |  |  |
| PxylGr42* | PxylGr42 |  |  |
| PxylGr43* | PxylGr43 |  |  |
| PxylGr44* | PxylGr44 |  |  |
| PxylGr45* | PxylGr45 |  |  |
| PxylGr46* | PxylGr46 |  |  |
| PxylGr47* | PxylGr47 |  |  |
| PxylGr48* | PxylGr48 |  |  |
| PxylGr49* | PxylGr49 |  |  |
| PxylGr50* | PxylGr50 |  |  |
| PxylGr51* | PxylGr51 |  |  |
| PxylGr52* | PxylGr52 |  |  |
| PxylGr53* | PxylGr53 |  |  |
| PxylGr54* | PxylGr54 |  |  |
| PxylGr55* | PxylGr55 |  |  |
| PxylGr56* | PxylGr56 |  |  |
| PxylGr57* | PxylGr57 |  |  |
| PxylGr58* | PxylGr58 |  |  |
| PxylGr59* | PxylGr59 |  |  |
| PxylGr60* | PxylGr60 |  |  |
| PxylGr61* | PxylGr61 |  |  |
| PxylGr62* | PxylGr62 |  |  |
| PxylGr63* | PxylGr63 |  |  |
| PxylGr64* | PxylGr64 |  |  |
| PxylGr65* | PxylGr65 |  |  |
| PxylGr66* | PxylGr66 |  |  |
| PxylGr67* | PxylGr67 |  |  |
| PxylGr68* | PxylGr68 |  |  |
| PxylGr69* | PxylGr69 |  |  |
| PxylGR1<sup>#</sup> | (delete) |  | Partial sequence of PxylGr75 |
| PxylGR2<sup>#</sup> | (delete) |  | Partial sequence of PxylGr34 |
| PxylGR3<sup>#</sup> | PxylGr72 |  |  |
| PxylGR4<sup>#</sup> | PxylGr73 |  |  |
| PxylGR5<sup>#</sup> | (delete) |  | Partial sequence of Gr77 |
| PxylGR6<sup>#</sup> | PxylGr75 |  |  |
| PxylGR7<sup>#</sup> | (delete) |  | Partial sequence of Gr77 |
| PxylGr64f-like<sup>※</sup> | PxylGr77 | XP_011560052.1 |  |
| PxylGr64f-like<sup>※</sup> | PxylGr78 | XP_011560051.1 |  |
| PxylGr28b<sup>※</sup> | PxylGr79 | XP_011565110.1 |  |

\*: Named by Engsontia et al., 2014; <sup>#</sup>: Named by Yang et al., 2017; <sup>※</sup>: Acquired from the GenBank

**Table S2.**
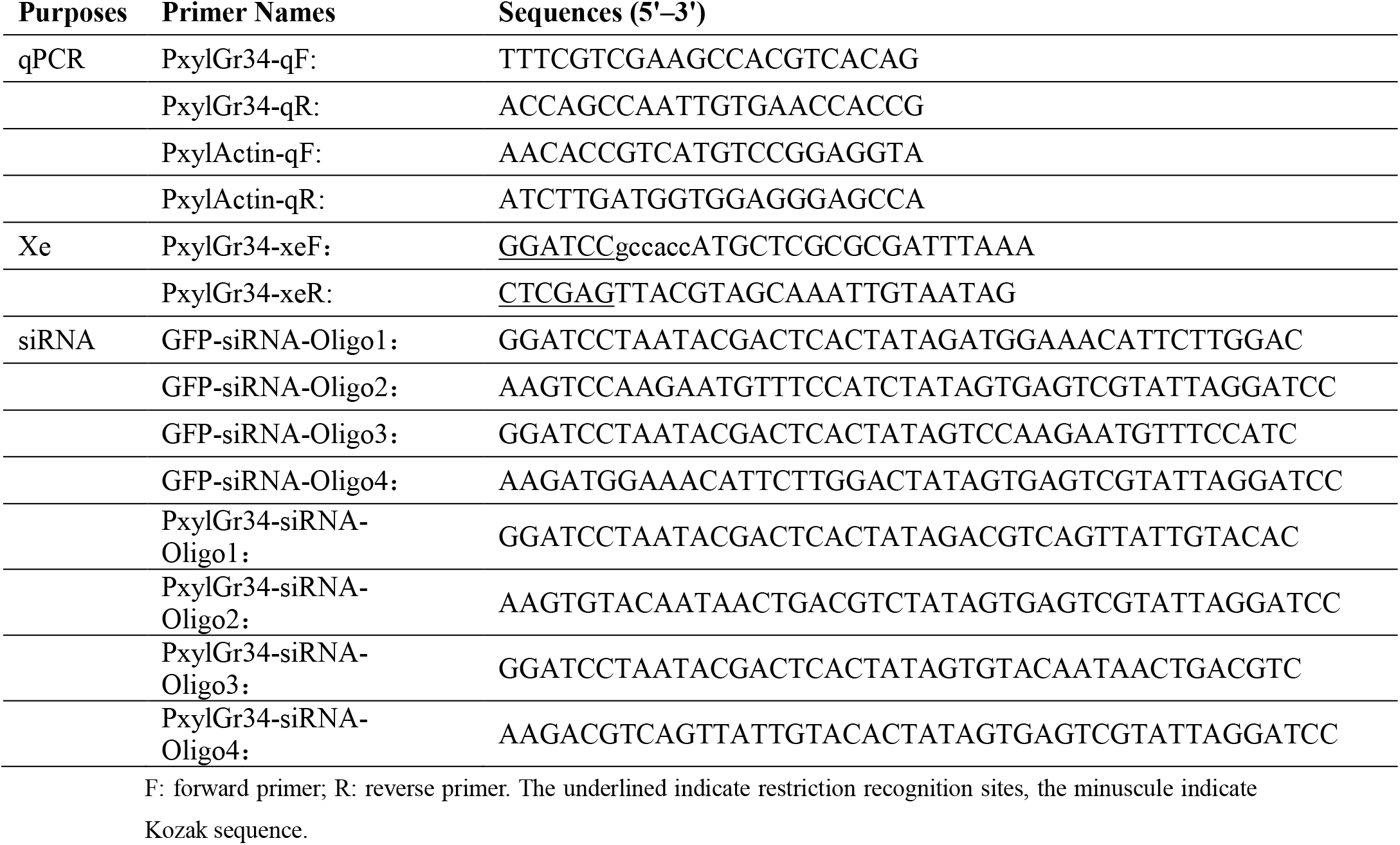
Primers Used for qPCR, *Xenopus* Oocyte Expression (Xe), and siRNA Synthesis.

